# Translesion synthesis protein ImuA from *Mycolicibacterium smegmatis* is a hexameric helicase-nuclease

**DOI:** 10.64898/2026.08.14.744951

**Authors:** Shehreen H. Khan, Harman S. Dev, Monica M. Warner, Dana J. Sowa, Kristi L. Lichimo, Samantha Reeve, Sara N. Andres

**Affiliations:** Biochemistry and Biomedical Sciences, McMaster University, Hamilton, Ontario, L8S 4K1, Canada; Michael G. DeGroote Institute for Infectious Disease Research, McMaster University, Hamilton, Ontario, L8S 4L8, Canada

## Abstract

Translesion DNA synthesis (TLS) enables DNA replication across damaged DNA and promotes stress-induced mutagenesis that contributes to antibiotic resistance in bacteria. The conserved ImuABC mutasome is essential for TLS in many bacterial species, yet the molecular function of its accessory protein, ImuA, has remained elusive. Here we show that *Mycolicibacterium smegmatis* ImuA assembles into a hexameric complex, likely arranged as a dimer of trimers, with dual enzymatic activities that reshape current models of its role in DNA damage tolerance. We show that ImuA functions as an ATP-dependent helicase that preferentially unwinds DNA substrates containing single-stranded DNA overhangs and identify amino acids required for both hexamer formation and helicase activity. Unexpectedly, ImuA also possesses ATP-independent 5′ exonuclease activity, selectively processing ssDNA substrates with free 5′ ends. We show a basic patch on the N-terminus is essential for stabilizing both the nuclease motif and oligomerization. Together, these findings identify ImuA as an active DNA-processing enzyme rather than a passive accessory factor and establish oligomerization as a prerequisite for its function. Our work provides a mechanistic framework for understanding how ImuA may function within the ImuABC mutasome to coordinate DNA processing during translesion synthesis.

## INTRODUCTION

Bacterial cells are continually exposed to DNA damaging agents arising from endogenous metabolic processes and environmental stressors. Reactive oxygen species, ultraviolet radiation, chemical mutagens, and certain antibiotics can produce backbone breaks, base modifications, crosslinks, and bulky adducts that distort the helix and impede the replication machinery.^1,2^ If left unrepaired, these lesions stall replication forks and threaten cell viability. Cells have therefore evolved damage-tolerance pathways that detect, repair, or bypass DNA lesions. One such mechanism is translesion DNA synthesis (TLS), which enables replication past lesions that stall the normal replicative polymerase, Pol III. TLS is inherently mutagenic, because the specialized polymerases that mediate lesion bypass have reduced fidelity and lack robust proofreading activity.^1–3^ This allows cell survival under genotoxic stress at the cost of an elevated mutation rate.

In approximately one-third of bacteria, including World Health Organization priority pathogens such as *Mycobacterium tuberculosis* and *Pseudomonas aeruginosa*,^4^ TLS is mediated by the ImuABC mutasome, a complex composed of ImuA, ImuB, and the specialized polymerase ImuC, also known as DnaE2, which belongs to the C-family of DNA polymerases.^5,6,8^ Genetic studies across diverse bacterial species have shown that the ImuABC-driven TLS pathway causes damage-induced mutagenesis, and that loss of any mutasome components reduces damage-induced mutation rates but impairs survival under genotoxic stress.^5,7–11^ In *P. aeruginosa*, knockouts of *imuB* or *imuC* yield substantially fewer ciprofloxacin-resistant mutants than wild-type.^10^ ImuC, the error-prone polymerase, is the principal mediator of DNA damage-induced mutagenesis in *M. tuberculosis*, where it promotes *in vivo* survival and the emergence of drug resistance during infection.^11^ The same dependence holds in *Mycolicibacterium smegmatis*, where ultraviolet irradiation and hydrogen peroxide increase the frequency of rifampicin-resistant mutants 108- and 16-fold in an ImuC-dependent manner.^9^ TLS therefore, acts not only as a survival mechanism, but as an evolutionary pathway that accelerates the acquisition of antimicrobial resistance, making the mutasome components attractive targets for therapeutics aimed at limiting bacterial adaptation.

Despite the importance of TLS, the functions of the accessory proteins ImuA and ImuB remain poorly understood. Sequence analysis indicates that ImuB is related to Y-family DNA polymerases but lacks the catalytic acidic residues required for polymerase activity, suggesting that it functions as an inactive polymerase homolog.^12^ ImuB also contains a conserved β-clamp binding motif that allows it to associate with the replication machinery and localize to sites of DNA damage.^13^ Biochemical and genetic studies suggest that ImuB interacts with both ImuC and ImuA, and our prior work has shown that *Myxococcus xanthus* ImuB forms a stable trimer in solution.^13–15^ Together, these observations are consistent with a role for ImuB as a scaffold that recruits TLS components to stalled replication forks.

The function of ImuA is considerably more elusive. ImuA proteins share limited sequence identity (∼30%) with the recombinase RecA but retain the overall structural fold based on Alphafold3 modelling predictions.^15–17^ RecA carries out homologous recombination, a DNA double-strand repair pathway, through ATP-dependent nucleoprotein filament formation on single-stranded DNA.^18^ Several motifs required for RecA polymerization are not conserved in ImuA homologs which argues that ImuA does not assemble into canonical RecA-like filaments.^6,15,19^ However, the filament is not the only assembly available to this fold as RecA shares a conserved nucleotide-binding core with ring-shaped hexameric helicases.^20,21^ Superfamily 4 and 5 helicases, which include DnaB, Rho, and bacteriophage T7 gp4, are hexameric rings built on the RecA fold, and RecA itself forms hexameric rings in the absence of DNA.^20–22^ In these rings the ATPase sites are composite and lie at the interfaces between adjacent subunits, so that one protomer binds the nucleotide while a neighbouring monomer contributes catalytic residues in trans.^22,23^ Ring assembly is therefore a prerequisite for nucleotide turnover, where oligomeric state of a RecA fold protein dictates function.

Given that ImuA is hypothesized to retain the RecA fold, we previously studied ImuA from *Myxococcus xanthus* and found conserved ATPase activity that was stimulated by DNA.^15^ Here we characterize ImuA from *Mycolicibacterium smegmatis*. ImuA forms a hexamer that is an ATP-dependent DNA helicase, unwinding duplex DNA substrates with either a 3′ or 5′ single-stranded overhang with comparable efficiency. The ImuA hexamer also has ATP-independent nuclease activity that requires a free 5′ end for engagement. Guided by an AlphaFold3 model showing a dimer of trimers arrangement, we mutated amino acids at the predicted ATP-binding site and amino acids that appear to stabilize this motif. These ImuA mutants result in ImuA trimers that lose helicase activity while retaining nuclease activity, indicating that the ATP-binding motif stabilizes the hexameric helicase-active form of ImuA. We also mutated amino acids that appear to stabilize a PDEEXK/Q nuclease motif. This mutant also formed a trimer and lacked nuclease activity on single-stranded DNA. These results suggest that ImuA may function as a DNA processing factor in TLS.

## MATERIALS AND METHODS

### Cloning

Codon optimized ImuA from *M. smegmatis* was ordered from Genscript and cloned into an expression plasmid through Gibson assembly (New England Biolabs) cloning^24^. Codon optimized expression plasmids for ImuA mutants R83A/S84A, D181A/Q183A, and K24A/K31A were ordered from Genscript. The ImuA R22A/R23A/K24A/K31A plasmid was made by site-directed mutagenesis (SDM)^25^ using oligonucleotide primers ordered from Integrated DNA Technologies (Supplemental Table 1). All plasmids were verified through whole-plasmid sequencing (Plasmidsaurus).

### Recombinant Protein Expression and Purification

Expression plasmids for all ImuA proteins were transformed by standard heat shock into *Escherichia coli* competent cells. Transformed cells were grown in Luria-Bertani broth and protein expression was induced with IPTG at low temperature for approximately 18 h. Bacterial cell pellets were resuspended in a Tris-based lysis buffer containing salt, a reducing agent, glycerol, mild detergents and lysozyme, supplemented with a protease inhibitor cocktail. Cells were lysed by sonication, and lysate was clarified by centrifugation.

Clarified lysate was purified by immobilized metal affinity chromatography (IMAC) using Ni-NTA resin and eluted with an imidazole gradient. Fractions containing ImuA were further purified by anion exchange chromatography over a salt gradient, followed by size-exclusion chromatography in a HEPES-based buffer containing salt and glycerol. Eluted protein was buffer exchanged and concentrated using centrifugal concentrators. Final purified protein fractions were assessed for purity by SDS-PAGE (Supplementary Fig. 1) and stored at −80 °C. Protein concentrations were determined by absorbance at 280 nm and are expressed for the monomeric state. Final purified protein fractions were visualized by SDS-PAGE to assess purity. SDS-PAGE of purified proteins is available in Supplementary Figure 1. Proteins were stored at −80°C. All proteins were quantified by absorbance at 280nm with the DS-11 microvolume spectrophotometer (Denovix) and all protein concentrations are expressed for the monomeric state.

### DNA substrate preparation

DNA substrates used in biochemical assays were synthesized by Integrated DNA Technologies. Oligonucleotide sequences are described in Supplementary Table 1. dsDNA substrates were annealed by mixing equimolar amounts of complementary strands, heating at 95 °C for 2 minutes and then cooled to 25°C over 45 minutes. DNA substrates were purified by size-exclusion chromatography (Superdex 200 Increase 10/300 GL column, Cytiva) equilibrated in buffer (10□mM Tris–HCl, pH□7.5, 500□mM NaCl, 2□mM EDTA pH□8.0). Purified DNA was pooled and concentrated using centrifugal micro-concentrators (Millipore) before final purification by ethanol precipitation.^26^

### Helicase Assays

Helicase assays were based on a previously published protocol, with modifications made to track the reaction over time.^27^ Briefly, 20uL reactions containing 10nM of a 6-FAM labelled dsDNA substrate (3’ overhang, 5’ overhang, blunt) (Supplementary Table 1) were incubated with 1.25uM of protein on ice for 5 minutes in helicase buffer (20mM Tris-HCl pH 8.0 and 5mM MgCl_2)_. Reactions were initiated with the addition of 2mM ATP and 500nM unlabelled trap DNA complementary to the unlabelled strand of the dsDNA substrate (Supplementary Table 1). Reactions were incubated in a water bath at 37°C and quenched at various time-points with 4uL of stop buffer (2% SDS, 200mM EDTA, 40% glycerol, 0.3% bromophenol blue). Reactions were loaded on 20% Native PAGE gels and ran for 45 minutes in 1X TBE before being imaged on an Amersham Typhoon imager (Cytiva) with excitation and emission wavelengths of 495nm and 520nm respectively for 6-FAM fluorescence. Gels were quantified using ImageJ. Fixed rectangular ROIs were placed over the duplex band, the displaced single-stranded band, and the degradation product region in each lane, and each species was background-corrected using a matched blank ROI. The degradation product regions were considered anywhere in the lane other than the duplex and single-stranded regions, since degraded product can be various sizes and this ensures all duplex signal loss is accounted for. Background-corrected intensities were summed to give the total fluorescent signal per lane, and each species was expressed as a percentage of that total, controlling for lane-to-lane differences in loading and signal recovery. Unwinding and degradation were expressed as the percentage of signal gained relative to a no-enzyme control, in which the DNA remained a fully intact duplex with no product formation. Degradation products were included in the total so that nuclease-dependent substrate loss was not misattributed to unwinding. Data was plotted and analyzed using Prism v. 10.4.2 (GraphPad).

### Nuclease Assays

Nuclease activity was tested on 6-FAM-labelled single-stranded DNA substrates (3’ labelled, 5’ labelled) (Supplement Table 1). For concentration-based assays, reactions (10 µL) were initiated by mixing 5uL aliquots of serial dilutions of ImuA with 5uL of 2x reaction mix (40 mM Tris-HCl pH 8.0, 10 mM MgCl□ 100 nM DNA substrate) containing no ATP or 4mM ATP. Reactions were incubated at 37 °C for 5 or 10 minutes and quenched with 10 uL of stop buffer (98% formamide, 10 mM EDTA). For time course assays, 70uL of 500 nM ImuA was mixed with 70uL of 2x reaction mix containing no ATP. The reaction was incubated at 37°C and 10 µL aliquots were withdrawn and quenched in 10 uL of stop buffer at various time points up to 10 minutes. No-enzyme controls received protein dilution buffer (50 mM HEPES pH 8.0, 400 mM NaCl, 10% glycerol) in place of protein. Quenched samples were heated at 95 °C for 10 min before loading on a 20% denaturing 3M UREA-PAGE gel and running in 1x TBE at 200 V for 1 h. Gels were imaged on an Amersham Typhoon imager (Cytiva) with excitation and emission wavelengths of 495nm and 520nm respectively for 6-FAM fluorescence and then quantified in ImageJ. Fixed rectangular ROIs of identical dimensions were placed over the full-length single-stranded substrate band and the degradation product region of each lane below, and each was background-corrected using a matched blank ROI. Background-corrected intensities were summed to give the total fluorescent signal per lane, and each species was expressed as a percentage of that total. Degradation was reported as the increase in the percentage of degradation product relative to the no-enzyme control. Data was plotted and analyzed using Prism v. 10.4.2 (GraphPad).

### Atomic force microscopy data collection and analysis

ImuA proteins were purified on a Superdex 200 Increase 10/300 GL column (Cytiva) using an AKTA Pure FPLC system in 50mM HEPES pH 8.0, 150mM NaCl to remove any aggregation contaminants and ensure protein purity prior to imaging. Purified protein was diluted to 100nM in AFM buffer (4mM HEPES pH 7.5, 23mM NaCl, 4mM MgCl2) and 20uL was deposited onto freshly cleaved mica (Ted Pella Inc) and allowed to incubate for 5 minutes. The mica was then rinsed with 1 mL of HPLC-grade water (Fisher Scientific) before residual water was blotted with filter paper and the mica fully dried with a stream of nitrogen gas. Images were captured in air in tapping mode using a Dimension Icon AFM (Bruker) with ScanAsyst-Air probes (Bruker). Nanoscope Analysis v.2.0 (Bruker) was used for image processing, including plane subtraction and flattening, as well as final image generation. Molecular volumes for all proteins were determined using Topostats.^28^ A standard curve was created to correlate calculated molecular volumes to known molecular weight using protein standard ovalbumin, conalbumin, aldolase, ferritin, and thyroglobulin (Supplemental Figure 2). Linear regression analysis of protein standard volume measurements was done using Prism v.10.2.2 (GraphPad) and mean molecular weight was calculated with the following standard curve equation: Volume= 2.56 x (Molecular Weight) +15.03. Statistical analyses were calculated with Prism v.10.2.2 (Graphpad).

## RESULTS

### ImuA forms a hexameric complex

The oligomeric state of ImuA is likely central to how it assembles within the mutasome, through association with ImuB.^13,14^ The *M. xanthus* homolog was reported to be predominantly monomeric in solution by mass photometry, so we asked whether the *M. smegmatis* homolog adopts the same state or instead self-associates into a higher-order species like its predicted homolog RecA.^15^ To address this, we performed single-particle atomic force microscopy (AFM) on SEC-purified ImuA (Figure 1A). Particles were distributed as a single population with no evidence of higher-order aggregation. A Gaussian fit to the volume distribution gave a mean molecular volume of 406.1 nm³ (95% CI 398.0 to 414.3 nm³), corresponding to a molecular weight of 152.8 kDa (Figure 1B). Given the theoretical monomer mass of ImuA at 24.07 kDa, this result was consistent with a hexameric assembly. Together, these data indicate that ImuA forms a stable hexameric complex, similar to the secondary oligomeric form of RecA.^20^

**Figure 1.**
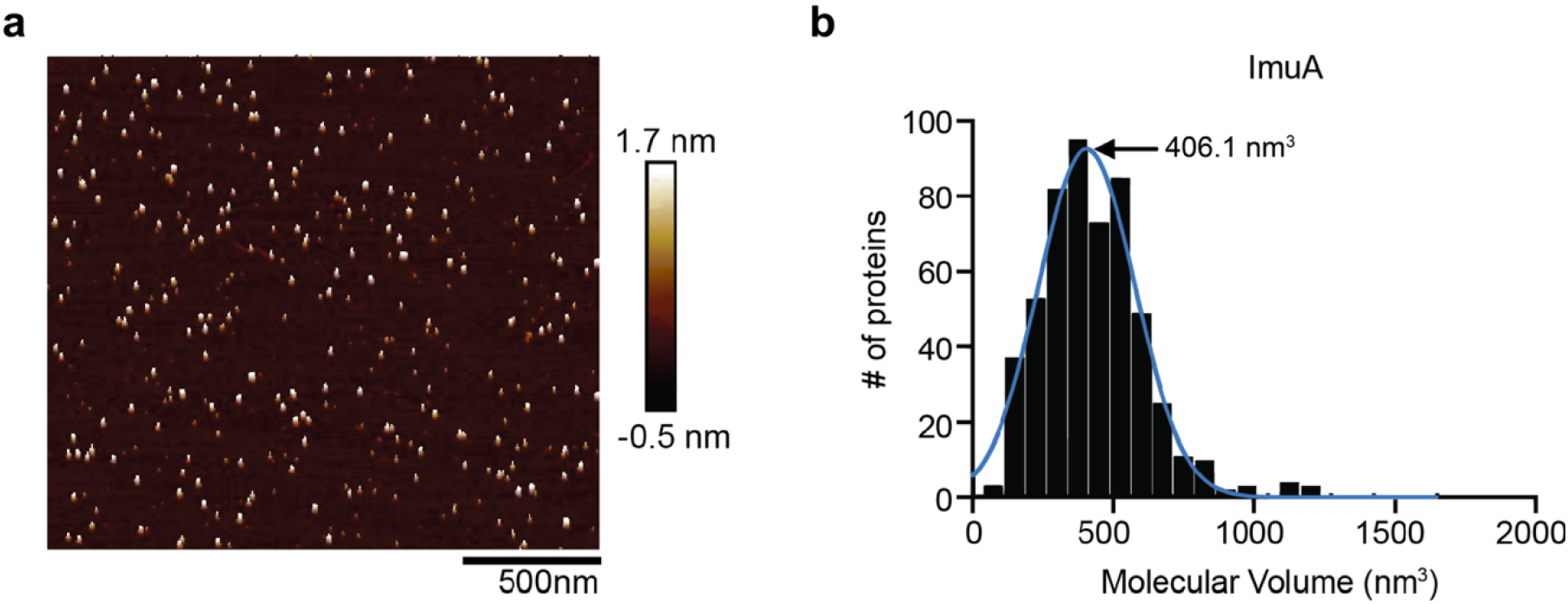
ImuA forms a hexamer. **a**, Representative AFM image of SEC-purified ImuA deposited on freshly cleaved mica. **b**, Molecular volume distribution of ImuA particles (n = 547). Blue line, Gaussian fit; mean molecular volume, 406.1 nm^3^ (95% CI 398.0–414.3 nm^3^).

### ImuA is a helicase that preferentially unwinds DNA substrates with single-stranded overhangs

RecA forms nucleoprotein filaments through an oligomerization interface that ImuA is not predicted to possess.^6^ However, the hexameric assembly observed by AFM (Figure 1) instead suggests an alternative RecA quaternary state in which the protein assembles into hexameric rings with structural homology to ring-shaped helicases.^20^ Given that our previous work showed ImuA from *M. xanthus* was a DNA-dependent ATPase, this prompted us to test directly whether ImuA is an ATP-dependent helicase.^15^ DNA unwinding was monitored using a gel-based strand displacement assay. On 24 bp duplex substrates bearing either a 5’ or 3’ 20-nt single-stranded DNA overhang, ImuA produced a loss of the intact duplex over time, where approximately 70% of the overhang substrates were depleted within 10 min (Figure 2A-D). In addition to the displaced single strand of DNA, we also observed faster-migrating degradation products, indicative of a potential nuclease activity, which is explored in greater detail below. This nuclease activity resulted in the single-stranded product to not accumulate stoichiometrically with substrate loss at later timepoints. We were unable to identify a separation of function mutant for helicase and nuclease activity, so dsDNA substrate loss was quantified alongside accumulation of single-stranded DNA, attributable to helicase activity, and accumulation of degradation products, attributable to nuclease activity, relative to total DNA in the lane (Figure 2A,C,E). We also investigated whether ImuA could act on blunt-ended DNA. The 40bp dsDNA was not depleted after 10 min (Figure 2E-F), with no substantial gain of either ssDNA or degradation products. Together, these data indicate that ImuA requires a single-stranded region to efficiently unwind duplex DNA and cannot degrade blunt-ended substrates.

**Figure 2.**
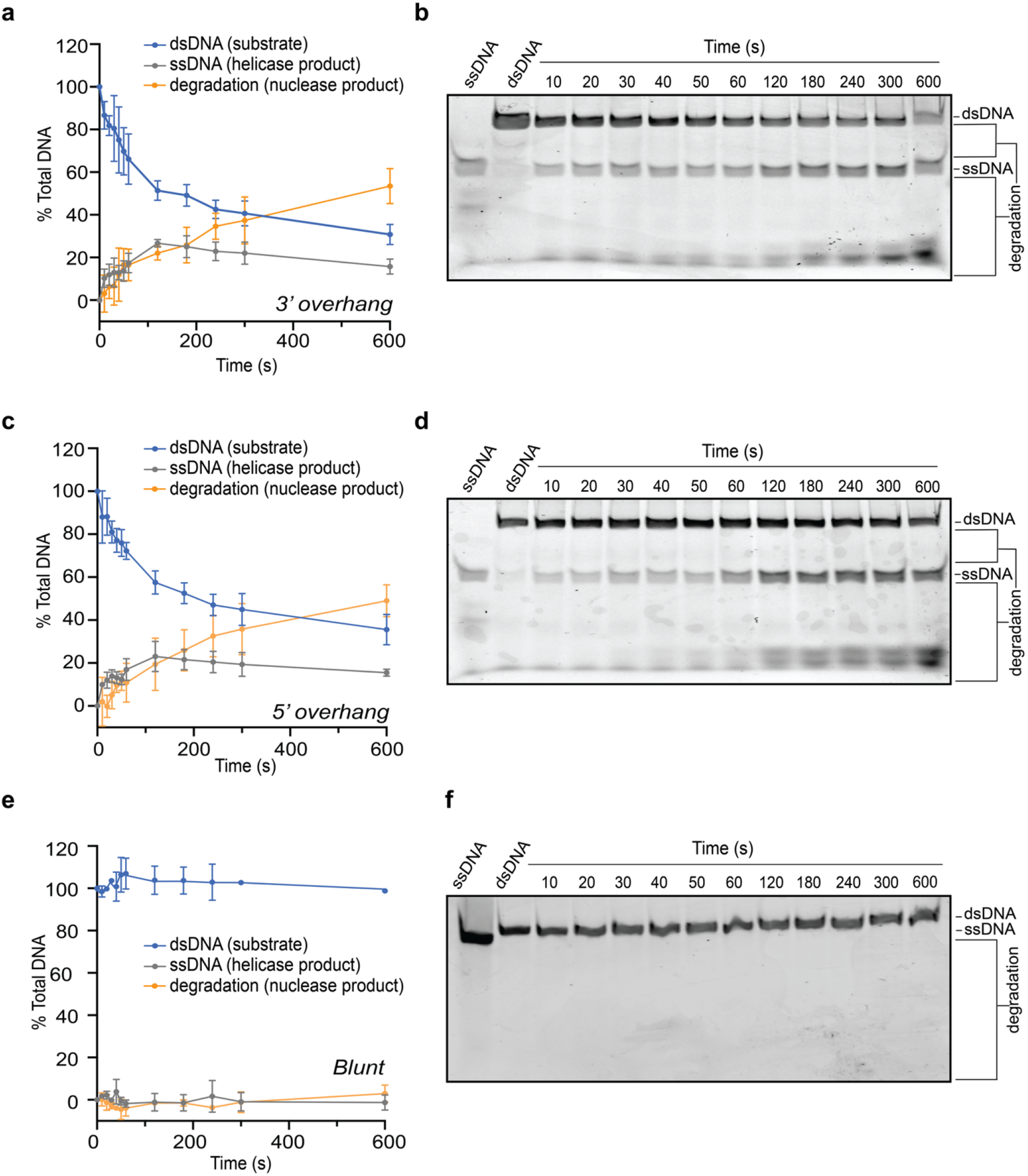
ImuA preferentially unwinds duplex DNA bearing single-stranded overhangs. Strand displacement assays were performed with 1.25 µM ImuA and 10 nM 6-FAM-labelled DNA substrate in the presence of 2 mM ATP and 5 mM MgCl_2_. Quantification of band intensities as a percentage of total lane intensity is plotted for **a**, 3′ overhang (n = 3), **c**, 5′ overhang (n = 3) and **e**, blunt (n = 2) substrates. Blue, intact duplex; grey, displaced single strand; orange, degradation products. Error bars, s.d. Representative gels are shown for the **b**, 3′ overhang, **d**, 5′ overhang and **f**, blunt substrates. The first two lanes of each gel contain single-stranded and double-stranded substrate markers, respectively, incubated in reaction conditions but in the absence of protein.

### ImuA R83/S84 and D181/Q183 are required for hexamerization and helicase activity but not nuclease activity

An AlphaFold3 (AF3) model of the ImuA hexamer was generated to examine how the subunits might assemble into a RecA-like helicase architecture and to identify residues that could support helicase and nuclease activity and mediate the hexameric interface.^20,29^ The most confident prediction was obtained when ATP and Mg^2+^ were added to the protein in stoichiometric ratios (Figure 3). Although the overall template modeling scores are low (ipTM = 0.2, pTM = 0.27), the pLDDT values indicated relatively higher confidence in the main protein fold compared to the extended N-terminal helices (Supplemental Figure 3). The model predicted a ring architecture organised as a symmetrically arranged dimer of trimers (Figure 3A-C).

**Figure 3.**
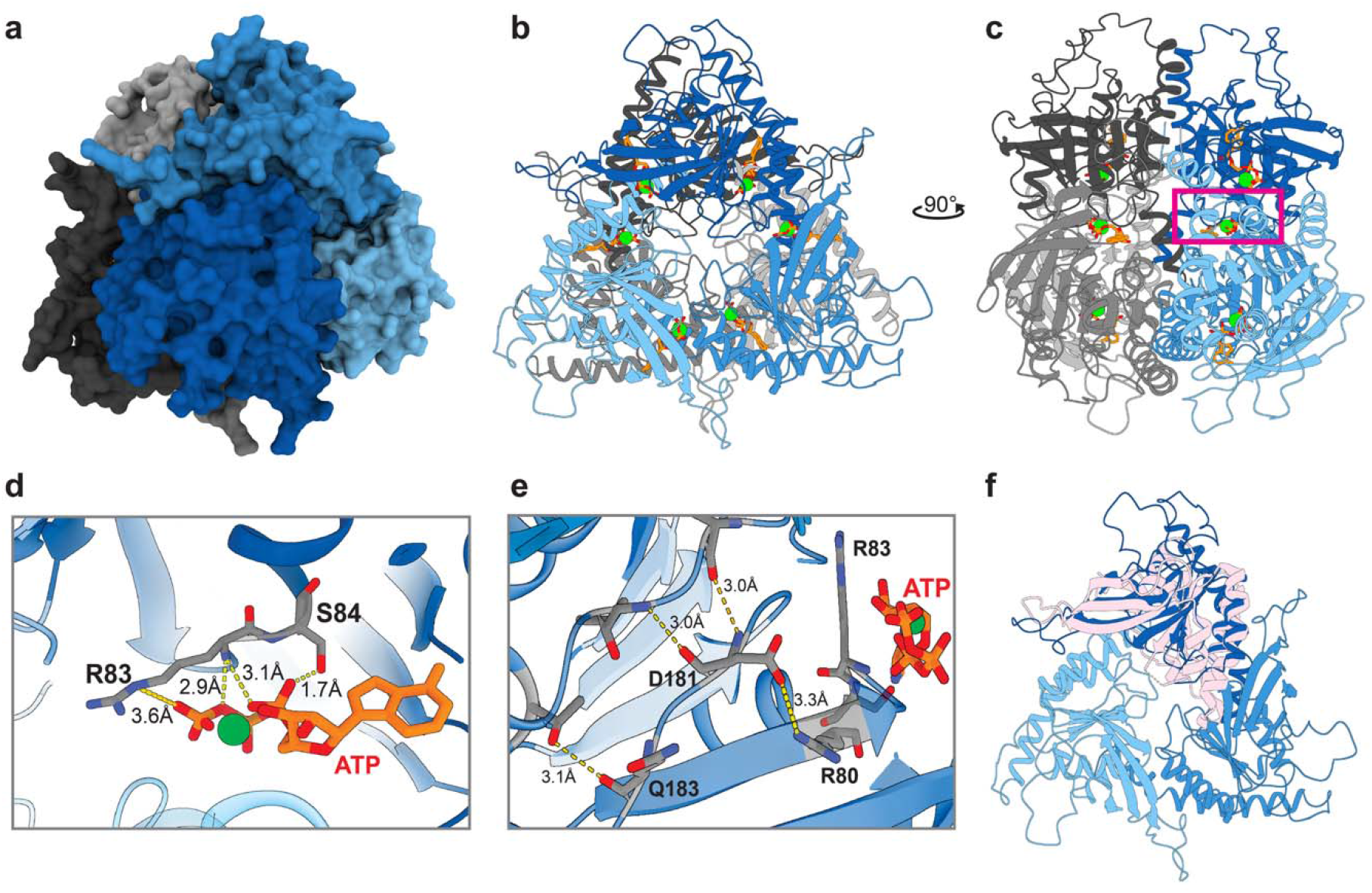
AlphaFold3 model of the ImuA hexamer reveals a dimer-of-trimers architecture with ATP bound between subunit interfaces. **a**, Surface model of AF3 prediction for the ImuA hexamer arranged as a dimer of trimers. Trimers are coloured blue and grey, with individual monomers in shades of blue or grey **b,** Secondary-structure representation of ImuA hexamer oriented facing a trimeric interface with ATP (orange) and Mg^2+^ (green spheres); **c**, Hexamer rotated 90° relative to **b,** oriented facing the dimeric interface; boxed region enlarged in **d** and **e**. **d**, R83 and S84 form hydrogen bonds with ATP and the coordinated Mg^2+^ ion. **e**, D181 lies within hydrogen-bonding distance of R80 on the ATP-binding helix, and Q183 hydrogen bonds to stabilize the strand. **f**, Superposition of a trimeric interface from the ImuA hexamer model (blue) with *M. smegmatis* RecA (PDB 1UBC, pink), RMSD of 2.7. Only the RecA main fold, lacking the N and C-terminal oligomerization domains, is shown for clarity.

ImuA R83 and S84 were located at the putative ATP-binding site, with both side chains within hydrogen-bonding distance of the ATP phosphates and the coordinated Mg² (Figure 3D). These residues align with the ATP-binding site, R71/T72, identified for the *M. xanthus* homolog in our previous work (Supplemental Figure 4).^15^ D181 and Q183 appear to support this motif, as D181 lies within hydrogen-bonding distance of R80 on the helix carrying the predicted ATP-binding residues, and Q183 anchors this region (Figure 3E). Alignment of this model with the crystal structure of an *M. smegmatis* RecA monomer (PDB 1UBC) yielded an RMSD of 2.7 Å, where the RecA ATP-binding site also matched that of the ImuA model (Figure 3F).

To test the functional importance of these identified elements and the validity of the AF3 model, we created two double point-mutants: ImuA R83A/S84A and ImuA D181A/Q183A, both designed to destabilize ATP binding. AFM analysis showed that ImuA R83A/S84A had an average molecular volume of 214.0 nm³ (95% CI 203.5 to 225.2 nm³), corresponding to a molecular weight of 77.7 kDa, and D181A/Q183A had an average molecular volume of 192.2 nm³ (95% CI 170.5 to 216.9 nm³), corresponding to 69.5 kDa (Figure 4A-D). Both values are consistent with a trimer of ImuA, as they are approximately half the volume measured for wild-type ImuA, indicating that each mutation prevents association of the trimers to form a hexamer, but maintains a trimeric interface. As the model shows the ATP-binding motif is stabilized by contacts across multiple interfaces, including between the dimeric interface, loss of hexamer formation is a plausible outcome.

**Figure 4.**
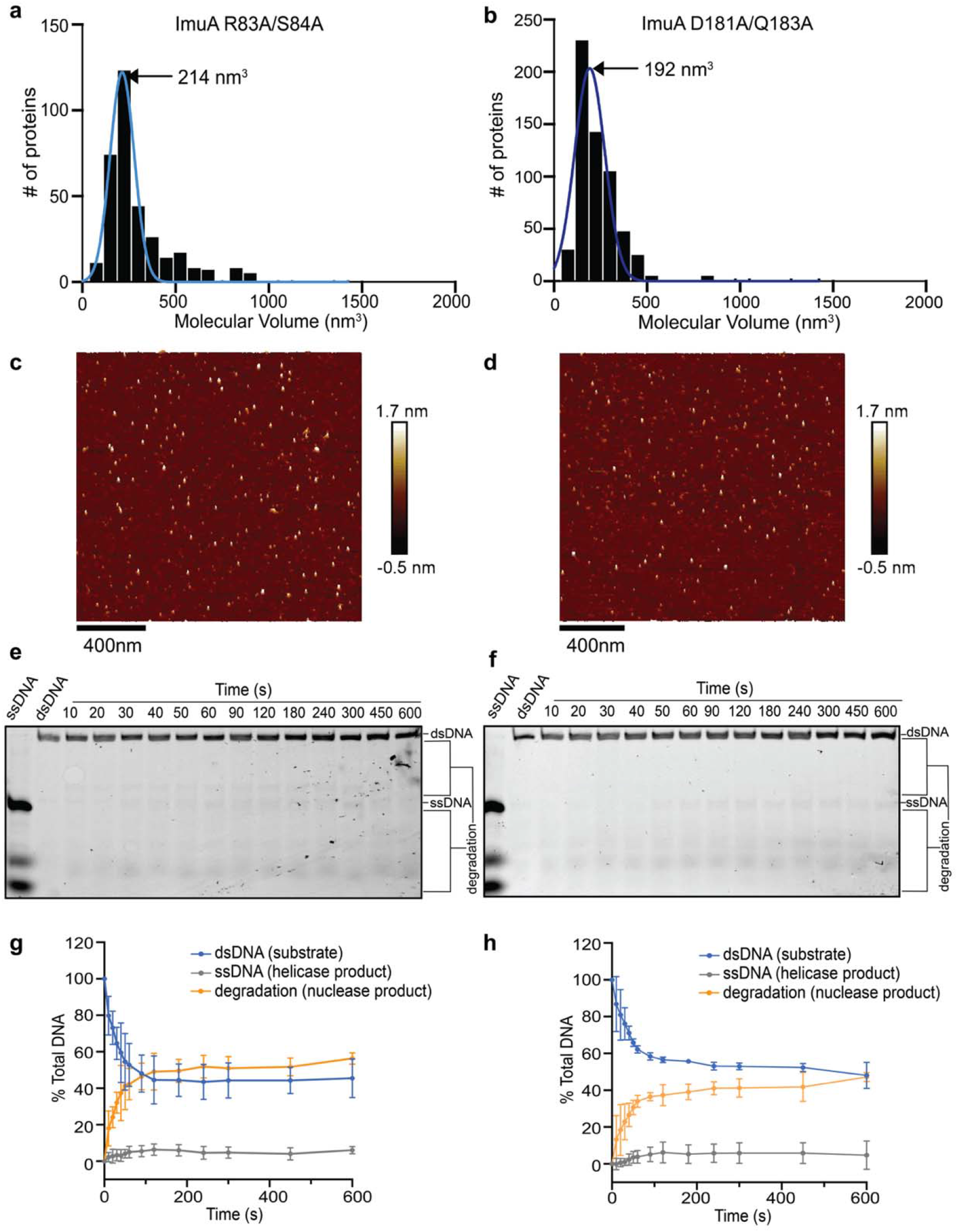
R83/S84 and D181/Q183 are required for hexamerization and helicase activity but not nuclease activity. **a**, **b**, Molecular volume distribution of ImuA R83A/S84A (**a**) and D181A/Q183A (**b**) particles. Blue line, Gaussian fit; mean molecular volume, 214.0 nm3 (95% CI 203.5–225.2 nm^3^) for R83A/S84A (n = 342) and 192.2 nm^3^ (95% CI 170.5–216.9 nm^3^) for D181A/Q183A (n = 240), corresponding to molecular weights of 77.7 kDa and 69.5 kDa, respectively, consistent with a trimeric assembly. **c**, **d**, Representative AFM images of SEC-purified ImuA R83A/S84A (**c**) and D181A/Q183A (**d**) deposited on freshly cleaved mica. **e**, **f**, Representative strand displacement gels for ImuA R83A/S84A (**e**) and D181A/Q183A (**f**) with 3′ overhang substrate. The first two lanes contain single-stranded and double-stranded substrate markers, respectively, incubated in reaction conditions but in the absence of protein. **g**, **h**, Quantification of band intensities as a percentage of total lane intensity for ImuA R83A/S84A (n=3) (**g**) and D181A/Q183A (n=3) (**h**). Blue, intact duplex; grey, displaced single strand (helicase product); orange, degradation products (nuclease product). Error bars, s.d.

Loss of the hexamer was also accompanied by loss of helicase activity. Where wild-type ImuA peaked at approximately 25% of single-stranded product accumulation before nuclease activity dampened signal, neither mutant produced substantial accumulation of the single-stranded product, remaining below 10% total DNA throughout the time course (Figure 4E-H). The intact duplex, however, was depleted to approximately 45% for both mutants, inversely correlating with the increase of degradation products. No initial delay of nuclease activity was observed with the ImuA mutants, as seen for the wildtype protein. Together, these results indicate that the R83/S84 and D181/Q183 interfaces are required for hexamerization and helicase activity, but dispensable for nuclease activity. This supports the AF3 model’s placement of these residues at the ATP-binding and oligomerization interface, with the observed ImuA mutants forming trimers consistent with the predicted dimer-of-trimers architecture.

### ImuA exhibits nuclease activity with apparent 5**′→**3**′** polarity

During our helicase studies, we observed degradation of the ssDNA reaction product along with helicase activity, suggesting ImuA possesses intrinsic nuclease activity. To characterize this activity directly, we tested concentration-dependent degradation of 3′-6-FAM-labelled ssDNA in the presence and absence of ATP (Figure 5A and B). ImuA degraded this substrate in a concentration-dependent manner, reaching approximately 80% degradation at 1 µM ImuA. Comparable degradation was observed with and without ATP, indicating that nuclease activity does not require ATP. To determine the directionality of this activity, ImuA was incubated with ssDNA substrates labelled with 6-FAM at either the 3′ or 5′ terminus (Figure 5C and D). Robust degradation was observed only for the 3′-6-FAM-labelled substrate, whereas the 5′-6-FAM-labelled substrate remained largely intact, indicating that ImuA requires a free 5′ end to initiate degradation, consistent with 5′→3′ exonuclease activity.

**Figure 5.**
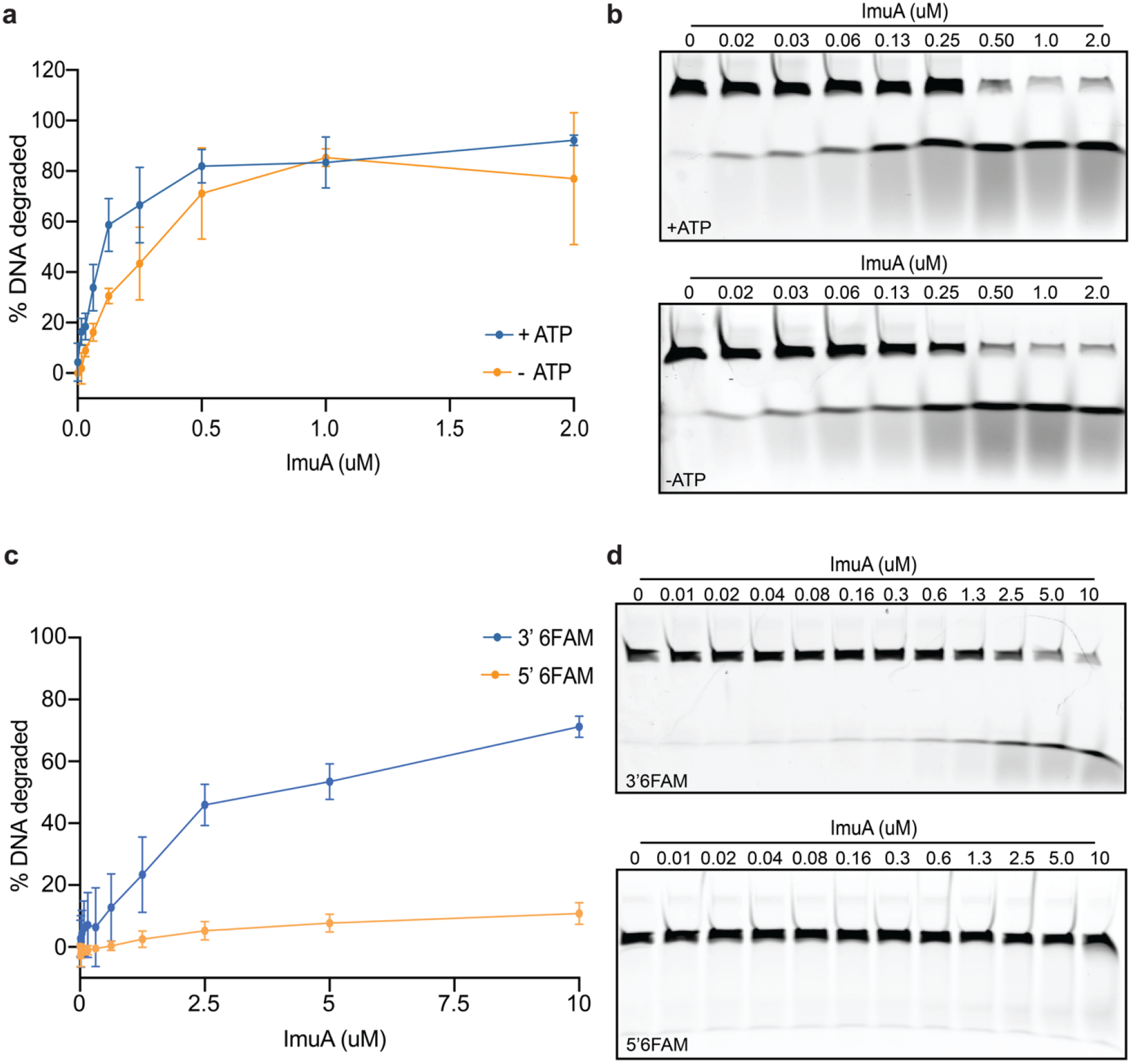
ImuA exhibits nuclease activity with apparent 5′→3′ polarity. Reactions contained 50 nM 6FAM-labelled ssDNA substrate in 20 mM Tris-HCl pH 8.0 and 5 mM MgCl2, with or without 2 mM ATP. **a**, Concentration-dependent degradation of 3′6FAM-labelled ssDNA by ImuA in the presence and absence of ATP, incubated at 37 °C for 10 min (n = 3). **b**, Representative gels corresponding to **a**, with ImuA titrated in the presence (top) and absence (bottom) of ATP. **c**, Degradation of ssDNA labelled with 6FAM at the 3′ or 5′ terminus by increasing concentrations of ImuA, incubated at 37 °C for 5 min (n = 3). **d**, Representative gels corresponding to **c**, with ImuA titrated against 3′6FAM-labelled (top) and 5′6FAM-labelled (bottom) ssDNA. Error bars, s.d.

### An N-terminal basic patch is required for nuclease activity on ssDNA but not on dsDNA

ImuA’s nuclease activity was surprising, as no nuclease motifs have previously been proposed for this protein. We therefore used the model to search for a candidate nuclease motif. In the AF3 hexamer model, the ImuA N-terminus forms an extended helix that forms the trimer-trimer interface, placing a cluster of basic residues, R22, R23, K24 and K31, at its base (Figure 6 A-B). These residues are wedged between adjacent monomers and appear to hold the long flexible N-termini in place. R11 is stabilized by this basic cluster, and reaches into the ATP-binding site of an adjacent monomer, potentially acting as an arginine finger for the nucleotide-binding motif (Figure 6C).^30,31^ Also stabilized by this basic cluster are residues Q17 and E19 of the N-terminus, placing them near P127, D128 and E135 of the same monomer forming a PDEEXK/Q-like nuclease motif (Figure 6B,D), suggesting this could be a plausible nuclease motif.^32–24^ To test the importance of this basic cluster on nuclease activity, we generated a charge-neutralizing mutant, ImuA R22A/R23A/K24A/K31A (ImuA RRKK).

**Figure 6.**
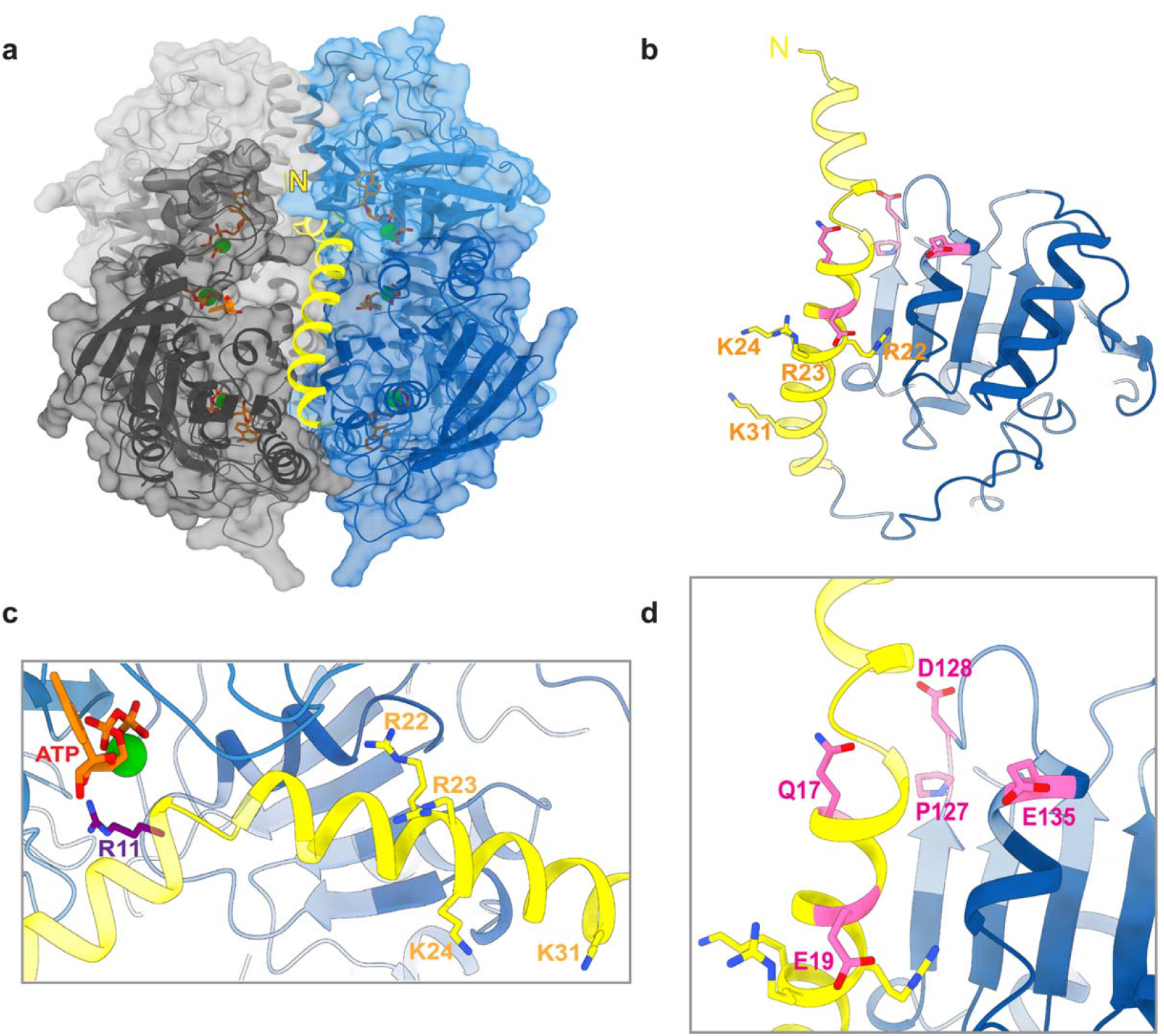
The N-terminal basic patch stabilized the trimer-trimer interface and positioned a candidate nuclease motif. **a** Surface representation of AF3 ImuA hexamer model with the N-terminal helix (yellow) shown at the trimer-trimer interface. **b.** Secondary structure of ImuA monomer seen in **a**, the basic patch residues R22, R23, K24 and K31 sits at the base of the N-terminal helix (yellow). **c.** R11 (purple) reaches into the ATP-binding pocket of the adjacent monomer near the bound ATP (orange/green), consistent with a possible arginine finger role, while R22, R23, K24 and K31 (yellow) anchor the N-terminal helix at the monomer interface. **d.** Q17 and E19 (pink) on the N-terminal helix are positioned near P127, D128 and E135 (pink), forming a PDEEXK-like nuclease motif.

AFM analysis showed that the ImuA RRKK mutant had a mean molecular volume of approximately 202 nm³ (95% CI 179.8 to 229.7 nm³) (Figure 7A, B), corresponding to a molecular weight of 73.2 kDa and consistent with a trimeric assembly similar to that observed for the R83A/S84A and D181A/Q183A mutants. Where wild-type ImuA degraded a ssDNA substrate ∼40%, ImuA RRKK showed no detectable degradation, consistent with our hypothesis that this basic patch stabilizes the predicted nuclease motif (Figure 7C-E). Unexpectedly, when ImuA RRKK was assayed for helicase activity against dsDNA bearing a ssDNA overhang, both unwinding and nuclease activity were observed at levels comparable to wild-type (Figure 6F, G). These results suggest a model in which dsDNA engagement stabilizes the ImuA N-terminus to support both ATP hydrolysis and correct positioning of the nuclease motif, offering a possible explanation for the recovery of nuclease activity by ImuA RRKK on dsDNA-containing substrates.

**Figure 7.**
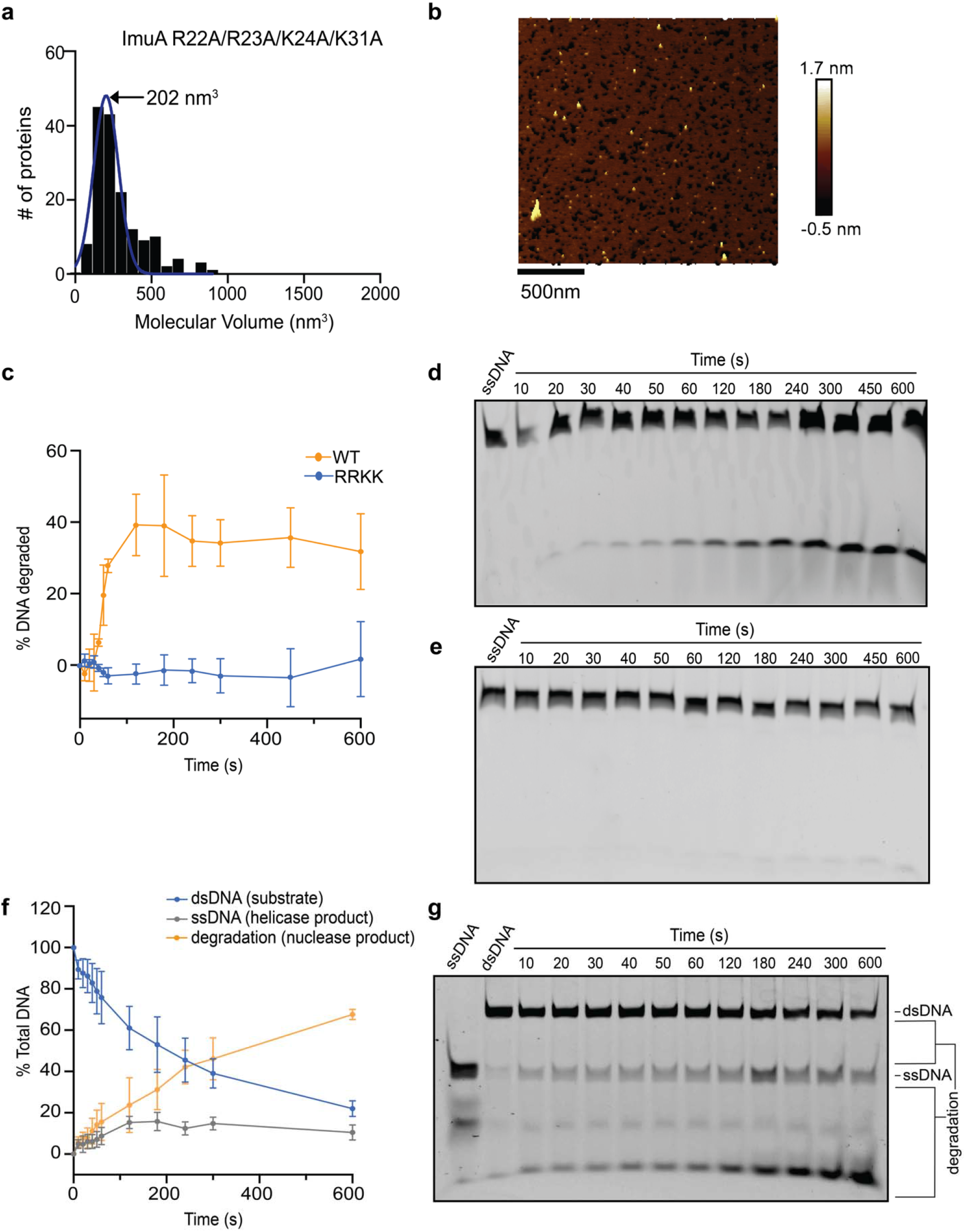
The N-terminal basic patch is required for nuclease activity on ssDNA but not on dsDNA-containing substrates. **a** AFM-based molecular volume distribution of ImuA RRKK (R22A/R23A/K24A/K31A), with a mean volume of ∼202 nm³ consistent with a trimeric assembly (n = 159). **b** Representative AFM image of ImuA RRKK. **c** Quantification of ssDNA degradation over time for WT ImuA and RRKK (n=3). **d** Representative gel showing WT ImuA-mediated degradation of 3′6FAM-labelled ssDNA over time. **e** Representative gel showing RRKK activity on 3′-6FAM-labelled ssDNA over time, with no detectable degradation. **f** Quantification of RRKK helicase and nuclease activity over time on a dsDNA substrate bearing a single-stranded overhang, showing loss of dsDNA substrate, generation of ssDNA helicase product, and generation of degradation nuclease product (n=3). **g** Representative gel showing RRKK activity on the duplex substrate over time, with dsDNA, ssDNA and degradation products indicated.

## DISCUSSION

Our findings redefine ImuA from *M. smegmatis* as an active DNA-processing enzyme rather than a catalytically passive accessory protein within the ImuABC mutasome. We show that ImuA assembles into a stable hexamer, functions as an ATP-dependent DNA helicase, and additionally possesses 5′→3′ exonuclease activity. Together, these observations indicate that ImuA likely plays a more direct role in DNA metabolism during translesion synthesis than was previously appreciated.

The discovery that ImuA forms a hexamer is significant because it extends the current view of ImuA beyond a RecA-like protein lacking the conserved motifs required for RecA filament formation, revealing instead that ImuA can adopt a higher-order oligomeric state. Interestingly, we previously observed ImuA from *M. xanthus* as monomeric by mass photometry.^15^ However, these observations may reflect species or methodological differences. The two proteins differ substantially in cysteine content, and mass photometry is performed under reducing conditions at lower concentrations that may have destabilized higher-order assemblies. Whether oligomerization varies among bacterial species or is influenced by cellular redox conditions or protein concentration remains an interesting question for future investigation.

The ATP-dependent helicase activity reported here provides a functional explanation for previously unexplained observations regarding ImuA biology. Previously, we observed that ImuA homologs preferentially bind DNA substrates containing single-stranded regions and exhibit DNA-stimulated ATPase activity.^15^ Here we find that efficient DNA unwinding required the presence of a single-stranded DNA overhang, which is also consistent with the loading requirements of many ring-shaped helicases.^35^ Together, these findings support a working model of TLS in which ImuA loads onto exposed single-stranded DNA generated at stalled replication forks and uses ATP hydrolysis to remodel DNA structures encountered during translesion synthesis.

Electrostatic surface mapping of the ImuA hexamer model revealed a strongly electropositive central channel, consistent with a role in nucleic acid engagement, as observed with other ring helicases (Figure 8A).^36^ Closer examination of the channel collar, formed by the TDGDWQGP loop (residues 178-185) contributed by each protomer of a trimer, indicated that this electropositive surface converges on a narrow acidic gate (Figure 8B). D181 from each protomer projects inward to form a symmetric ring with pairwise distances of 6 Å, positioned beneath the wider W182-lined mouth of the channel (24 Å pairwise distance). These observations align with other ring helicases, such as eukaryotic CMG helicases, where the channel is dynamic and can change in width upon DNA binding, to permit or block access of ds- or ssDNA.^37^ Structural superposition with M. smegmatis RecA (PDB: 1UBC) positioned this loop against the L2 DNA-binding loop of RecA (NQLREKIGVMFGSPET, residues 195-210), a region that directly contacts DNA in canonical RecA-family recombinases.^20^ The ImuA loop is shorter, and the compact sequence protruding into the central channel contains the acidic D181 and bulky W182 residues, raising the possibility that this loop was repurposed through sequence divergence to confer a distinct mode of nucleic acid engagement (Figure 8B). This divergence may reflect functional specialization for the substrate requirements identified in this study, including the dependence of helicase activity on single-stranded overhangs and the 5′-end dependence of nuclease activity.

**Figure 8.**
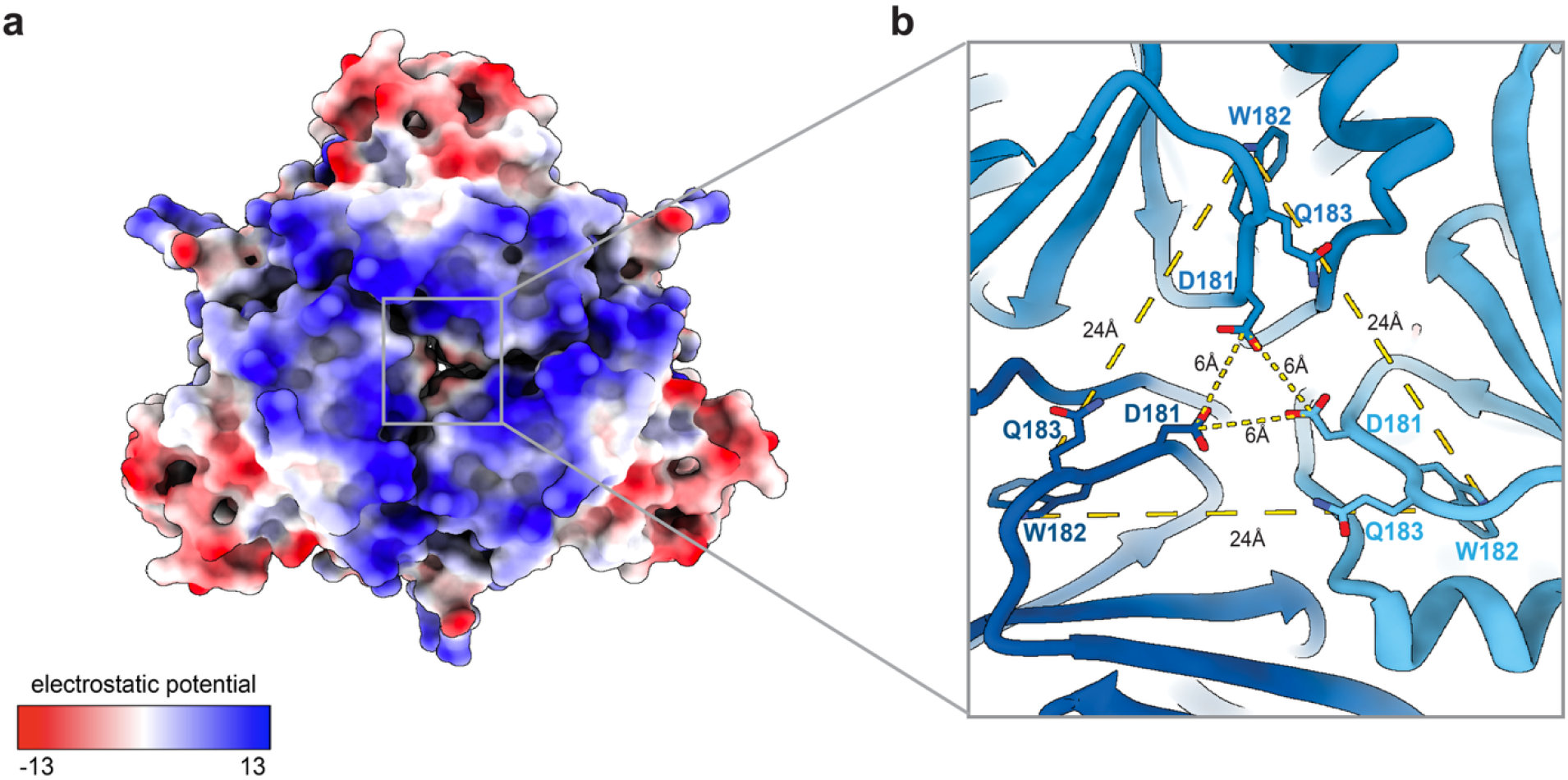
Electrostatics reveal an electropositive channel collar with an acidic gate. **a.** Surface representation of the ImuA hexamer colored by electrostatic potential from −13 to 13 kcal/(mol·e), revealing an electropositive central channel. **b.** Close-up of the channel collar formed by the TDGDWQGP loop (residues 178-185) across a trimer. W182 projects outward at the channel mouth with pairwise distances of 24 Å. D181 converges toward the channel axis with pairwise distances of 6 Å. Q183 flanks W182 at the periphery of each loop.

These biochemical activities also provide insight into previous studies showing that *M. xanthus* ImuA antagonizes homologous recombination through inhibition of RecA.^6^ In other bacteria, helicases such as UvrD and PcrA actively remove RecA nucleoprotein filaments from DNA through ATP-dependent translocation.^38,39^ Since previous findings have shown that ImuA can inhibit RecA directed homologous recombination,^6^ the discovery that ImuA is itself an ATP-dependent helicase suggests a plausible molecular mechanism by which this RecA paralog could regulate pathway choice at stalled replication forks in a similar manner to UvrD and PcrA. Rather than simply recruiting TLS components, ImuA may actively remodel DNA or protein-DNA complexes, removing RecA filaments, to favour lesion bypass over homologous recombination. Future studies are needed to confirm this model.

An unexpected finding of this study is that ImuA also possesses ATP-independent 5′→3′ exonuclease activity. The ATP independence of this activity distinguishes it from the helicase function, indicating that ImuA carries out at least two mechanistically distinct DNA-processing reactions. Multifunctional enzymes that combine helicase and nuclease activities are common throughout DNA replication and repair, where coordinated unwinding and strand processing enable efficient remodeling of complex DNA structures.^39^ For example, AddAB and RecBCD couple ATP-dependent DNA unwinding with nuclease activity to process double-strand break ends during homologous recombination, while eukaryotic DNA2 combines helicase and nuclease activities during Okazaki fragment maturation and DNA end resection.^40–43^ Likewise, the archaeal HerA-NurA system coordinates helicase-driven DNA translocation with nuclease-mediated processing during double-strand break repair.^44^ Although ImuA differs from these enzymes in both sequence and biological context, its ability to both unwind and degrade DNA suggests that it may similarly coordinate multiple DNA-processing activities during translesion synthesis. Whether these two activities act sequentially on the same DNA substrate, are independently regulated, or require interaction with ImuB or ImuC remains an important question for future work.

Building on this, mutagenesis data pointed to a structural basis for nuclease activity. Deletion of the N-terminal basic patch (R22A/R23A/K24A/K31A) abolished ssDNA degradation and disrupted hexameric assembly, indicating this patch is required for both oligomerization and nuclease function. The AF3 model offered a plausible explanation: the basic patch anchored the N-terminal helix at the trimer-trimer interface, positioning Q17 and E19 near P127, D128 and E135 to form a candidate PD-(D/E)XK-like motif, while the same helix placed R11 within the ATP-binding pocket of the adjacent monomer as a putative arginine finger. This proximity suggested the two motifs shared a common structural element, coupling ATP engagement to nuclease site positioning. Consistent with this, RRKK retained helicase and nuclease activity comparable to wild-type on a dsDNA substrate with a single-stranded overhang, indicating duplex engagement could stabilize the N-terminal helix independently of the basic patch and reposition the nuclease motif. This raised the possibility that ATP/DNA binding indirectly supports nuclease motif positioning through stabilization of the N-terminal helix, linking the ATPase and nuclease activities of ImuA in the context of duplex-containing substrates during translesion synthesis.

Overall, this work substantially expands the functional repertoire of ImuA and supports an evolving model of the ImuABC mutasome in which ImuA serves as an active catalytic component rather than a passive accessory factor. By combining ATP-dependent helicase activity with ATP-independent exonuclease activity within a defined oligomeric assembly, ImuA appears well positioned to remodel DNA substrates generated during translesion synthesis and coordinate mutasome function at stalled replication forks. Given the central role of ImuABC in damage-induced mutagenesis and the evolution of antimicrobial resistance, defining the molecular mechanisms that regulate ImuA activity may ultimately provide new opportunities to inhibit bacterial adaptation without directly targeting essential DNA replication.

## ACKNOWLEDGEMENTS

We acknowledge the Centre for Microbial Chemical Biology for instrumentation used in gel imaging. We also acknowledge the Centre for Advanced Light Microscopy at McMaster University for instrumentation used in atomic force microscopy.

## AUTHOR CONTRIBUTIONS

S.H.K. undertook the majority of the experimental design, data acquisition, analyses and writing of the manuscript. H.S.D. assisted with protein purification and helicase studies. M.M.W assisted with atomic force microscopy studies. D.J.S, K.L.L, and S.R. assisted with cloning of proteins used in this study. S.N.A. supervised project, provided funding and resources, co-wrote and edited the paper alongside S.H.K.

## DATA AVAILABILITY

The raw numbers for charts and graph, raw gel images and raw AFM images will be available at https://borealisdata.ca/dataset.xhtml?persistentId=doi:10.5683/SP4/LNGKUG. The Alphafold3 model will be available at https://modelarchive.org/doi/10.5452/ma-ngfij.

