## Supplemental Data for "Translesion synthesis protein ImuA from *Mycolicibacterium smegmatis* is a hexameric helicase-nuclease"


**Supplemental Table 1.** Oligonucleotide sequences of DNA substrates used in methodology.

| Substrate | Sequence | Use |
|---|---|---|
| ImuA KKRR FWD primer | 5’ CACCTGGCTGCTGCCATGGCATCCGTGTCTGCAGCTGTGGGCAGCGCG 3’ | SDM cloning |
| ImuA KKRR REV primer | 5’ CGGATGCCATGGCAGCAGCCAGGTGCTCAACTTGTTCCGCTCTCGTCAAGCGCCT 3’ | SDM cloning |
| 3’ overhang | 5’ACCTTCGTTGGTCGGCAGCAGGGCTTTTTTTTTTTTTTTTTTTT 3’<br>5’GCCCTGCTGCCGACCAACGAAGGT 6-FAM 3’ | Helicase assay |
| 5’ overhang | 5’TTTTTTTTTTTTTTTTTTTTACCTTCGTTGGTCGGCAGCAGGGC 3’<br>5’GCCCTGCTGCCGACCAACGAAGGT 6-FAM 3’ | Helicase assay |
| blunt | 5’ 6-FAM CAGGGTAAGTGTGGAGGTGTAGGGAAGGGAATGTTGTCTG 3’<br>5’ CAGACAACATTCCCTTCCCTACACCTCCACACTTACCCTG 3’ | Helicase assay |
| 3’6FAM ssDNA | 5’GCCCTGCTGCCGACCAACGAAGGT 6-FAM 3’ | Nuclease assay |
| 5’6FAM ssDNA | 5’ 6-FAM ATCGACTCTTGAGGACAGCA 3’ | Nuclease assay |

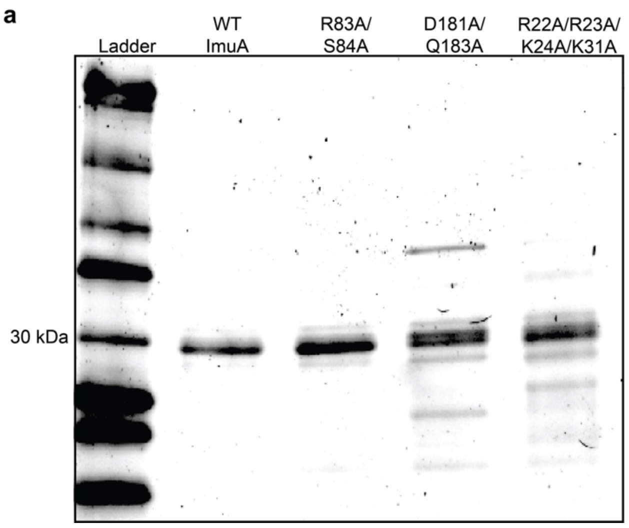


**Supplemental Figure 1. Purification of ImuA wild type and mutant proteins. a.** SDS-PAGE analysis of purified WT ImuA and the indicated mutants (R83A/S84A, D181A/Q183A and

R22A/R23A/K24A/K31A). All proteins migrated at the expected monomeric molecular weight of approximately 30 kDa.

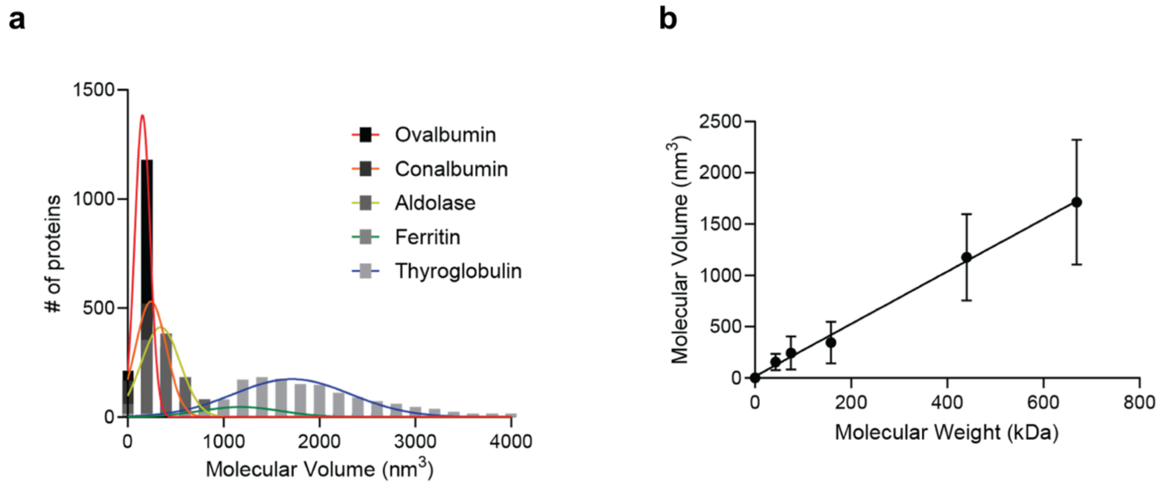


**Supplemental Figure 2. AFM standard curve. a.** Frequency distribution plot of volumes of protein standards (as listed in legend). Data represented as mean ± SD calculated from a nonlinear Gaussian fit. **b.** Standard curve of volumes corresponding to molecular weight (kDa) with equation of the line V = 2.556MW + 15.03 calculated with Prism v10.2.2. (GraphPad). Data plotted as mean ± SD. $R^2$=1.0.

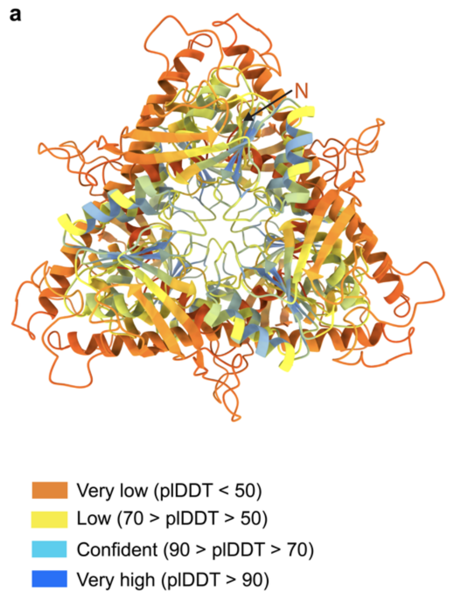


**Supplemental Figure 3. AlphaFold3 model of the ImuA hexamer colored by per-residue confidence (pLDDT). a**, Full-length hexamer model with extended N-termini included, viewed down the central pore (pTM= 0.2, ipTM= 0.27).

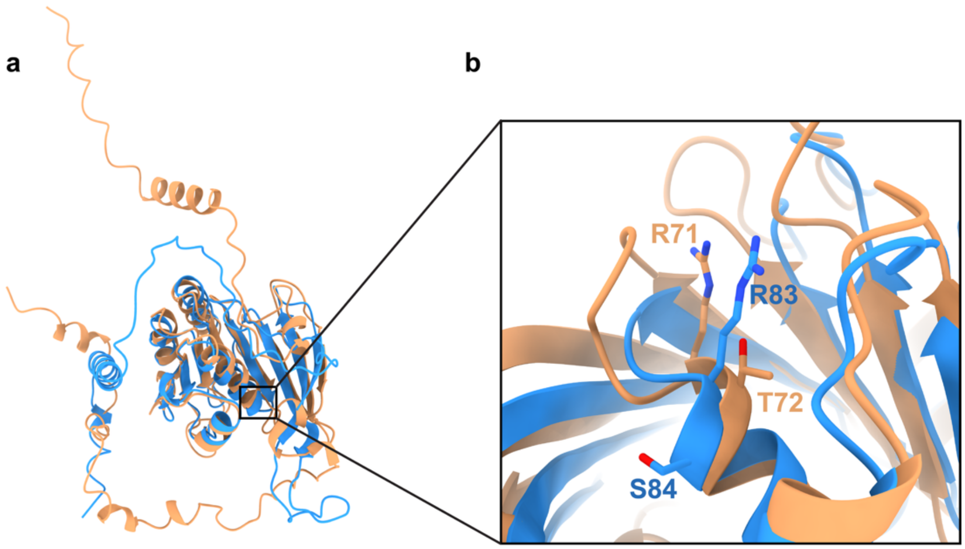


**Supplemental Figure 4. Alignment of AlphaFold3 models for the ImuA monomer from *M. smegmatis* and *M. xanthus*. a,** AF3 prediction of *M. smegmatis* ImuA (blue, pTM = 0.67) aligned with *M. xanthus* ImuA (tan, pTM = 0.69; RMSD = 1.33Å). **b,** Close-up of the previously identified ATP-site residues R71/T72 in *M. xanthus,*[15] corresponding to putative ATP-site residues R83/S84 in the *M. smegmatis* structure.
